# Genome-wide Perturbation Analysis Screening System for Exosomes-Related Genes Based on the CRISPR/Cas9 Platform

**DOI:** 10.1101/2023.09.07.556640

**Authors:** Lu Lu, Jianxin Yin, Ning Chen, Xiaofei Guo, Chunle Han, Miao Wang, Huanqing Du, Huifang Li, Xingang Pan, Mengya Gao, Na Wang, Dongli Qi, Jianhong Wang, Fengwei Dong, Tianshi Li, Xiaohu Ge

## Abstract

As reported previously, exosomes have significant impacts on the physiological and pathological state *in vivo*. Exosomes have been extensively studied as a drug delivery carrier and some exosomes drugs have been undergoing clinical research. The mechanisms of exosome production, transport, and secretion remain to be studied in depth. Thus, here, we developed a novel CRISPR-UMI-based gene screening system, with which we can reveal the exosomes biogenesis, transport, and uptake mechanisms on a genome-wide scale. This system consists of two parts: one is a gene knockout component; the other part is a unique molecular identifiers (UMI) labeling component that can label the exosomes produced by the knockout cells, in which each sgRNA corresponds to a specific UMI. In this way, by detecting the UMI loaded in the exosomes, we can trace the knockout gene. In this study, we first verified the function of each component using plasmids and lentiviruses respectively. Next, we simulated the infection of cells with lentiviral libraries using a single lentivirus to validate the functionality of the screening system. Finally, we constructed a CRISPR-UMI-based library targeting 15 genes (genes with clear effects on exosomes biogenesis) to further validate the reliability of the screening system. The development of this screening system is of indispensable importance for the in-depth study of the mechanisms of exosome production and secretion, as well as for the improvement of exosomes production and the advancement of exosomes industrialization.

## Background

Exosomes are lipid bilayer vesicles secreted by cells with a diameter between 40 and 150 nm separated from other extracellular vesicles. Exosomes were first discovered in 1983 in sheep reticulocytes [1]. Almost all cells are capable of secreting exosomes, and they are naturally present in various body fluids [2]. Exosomes are involved in intercellular communication and various biological processes through various substrates that they encapsulate or load, including RNA, DNA, protein, and lipids [3].

At first, exosomes were considered the “trash bag” of cells, allowing cells to get rid of some useless proteins. However, in recent years, studies have found that the “cargo” carried by exosomes has important biological significance [4]. Exosomes naturally carry the functions of mother cells. For example, exosomes released from mature dendritic cells (DC) can cause an adaptive immune response [5]. Based on this fact, Ying Tian et al. developed engineered Treg cell-derived exosomes for the treatment of ocular neovascular diseases [6]. Exosomes biogenesis can capture complex extracellular nuclear intracellular molecules. For example, at present, exosomal circulating RNAs (lncRNAs) have become an effective molecular marker for early cancer detection. Ling-Yun Lin et al. isolated exosomes from early gastric cancer patients and normal people and screened exosomal lncRNAs as potential biomarkers for early gastric cancer by lncRNAs sequencing and analysis [7]. Similarly, Liquid molecular Diagnostics developer Exosomes Diagnostics Inc. has promoted the analysis of exosomes-encapsulated circumscribed tumor DNAs (cfDNAs) as an effective *ex vivo* biopsy method for non-small cell lung cancer (NSCLC) [8]. Due to the heterogeneity of exosomes secreted by tumor cells, exosomes can provide a map for the diagnosis of the disease [9]. Exosomes also have good biocompatibility and the ability to target specific cells and transfer bioactive molecules [10]. Exosomes are naturally suitable for transporting proteins, mRNAs, miRNAs, various non-coding RNAs, mitochondrial DNAs, genomic DNAs, and so on. Zhenzhen Wang et al. used lung stem cell-derived exosomes loaded with RBD mRNAs to prepare a COVID-19 vaccine, which elicited an immune response [11]. Codiak loaded therapeutic agents such as Cyclic dinucleotide agonist [12] or ASO into exosomes and then delivered them to targeted cells or tissues, resulting in good preclinical results[13]. As a kind of nanovesicles, exosomes attract more and more people to apply them in drug delivery. Therefore, they not only have potential clinical applications in diagnosis but also in drug delivery. As drug delivery carriers, exosomes have irreplaceable advantages such as low immunogenicity, good biocompatibility, and biological stability[14], so they are of great commercial value. Large-scale production of clinical-grade exosomes is a necessary path to commercialize exosomes. The production of exosomes is variously limited by the exosomes biogenesis ability of their parent cells. So, improving the exosomes biogenesis of their parent cells will also contribute to the clinical translation of exosomes. A comprehensive understanding of the mechanisms of exosomes biogenesis is the premise of our engineering cells.

There have been several studies on exosomes biogenesis, which mainly focused on the endosome-sorting complex transport dependent (ESCRT-dependent) pathway[15]. Members of the ESCRT family of proteins have been shown to bind to a series of complexes (ESCRT-0, ESCRT-I, ESCRT-II, and ESCRT-III) on the MVB membrane to regulate the targeted transporting of molecular cargos into the ILVs and ILVs formation [16]. Until recently, several reports suggested exosomes could also be generated by non-ESCRT-dependent pathways. For example, members of the Rab family of small GTPases play promising roles in transferring vesicles between intracellular compartments, and exosomes releasing. nSMase2-ceramide pathway could drive ILVs formation and MVBs sorting in an ESCRT-independent mechanism. And there also are other ESCRT-independent mechanism pathways relative to proteins, for example, Caveolin□1, and some of the transmembrane proteins (CD9, CD63, CD81, etc.) [16]. However, it is likely that there are still other mechanisms related to exosomes biogenesis that need to be discovered. There were few studies on the systematic screening of the mechanism pathways involved in exosomes biogenesis. Here, we developed a novel CRISPR-UMI-based screening system, for the systematic screening of pathways related to exosomes biogenesis including exosomes formation, cargos loading, exosomes secretion, with minimal omissions.

CRISPR-UMI-based screening system cleverly combines UMI with gene knockout tools. UMI is a molecular barcode containing 10 bases. It is used to label exosomes produced by cells containing sgRNAs that uniquely correspond to UMI. This ingenious design allowed us to establish a correspondence between gene knockout and exosomes biogenesis. In addition, by delivering UMI-labeled exosomes to cells or tissues, we can monitor the uptake bias of exosomes in different cells or tissues and further track down the genes affecting the uptake bias of exosomes. Since the UMI diversity can fully cover the human genome-wide sgRNA library, we can use the CRISPR-UMI-based screening system to scan the whole genome and establish pathway relationships for certain needs, such as exosomes biogenesis, uptake, tissue bias, and so on. It is expected to draw a complete map of exosomes biogenesis from the perspective of systems biology. Moreover, there are no systematic studies reported on the mechanisms of exosomes biogenesis. This CRISPR-UMI-based system provides a powerful research tool for exploring the physiological and pathological mechanisms of exosomes.

## Result

### Working principle of the screening system

The working principle of the CRISPR-UMI-based screening system developed in this study is shown in Figure 1. As the figure shows, two diverse lentivirus particles successively infected 293T cells, the interaction of which ultimately induced the reduced expression of target genes and change of the labeled exosomes with the UMI corresponding to target genes concurrently. Specifically, we employed two different lentiviruses to make the screening system work. The BASP1-MCP-Cas9 lentivirus served as an endonuclease and exosomes-located RNA-binding protein provider. The sgRNA-UMI lentivirus library contained sgRNA sequences targeting different genes and the corresponding UMI-MS2 labels. The fusion of exosomes location protein (BASP1) and RNA-binding protein (MCP) can localize MCP into exosomes [19], thereby bringing its recognition sequence (UMI-MS2) into exosomes during exosomes biogenesis. These exosomes were then isolated and purified for total RNA extraction and NGS analysis.

**Figure1.**
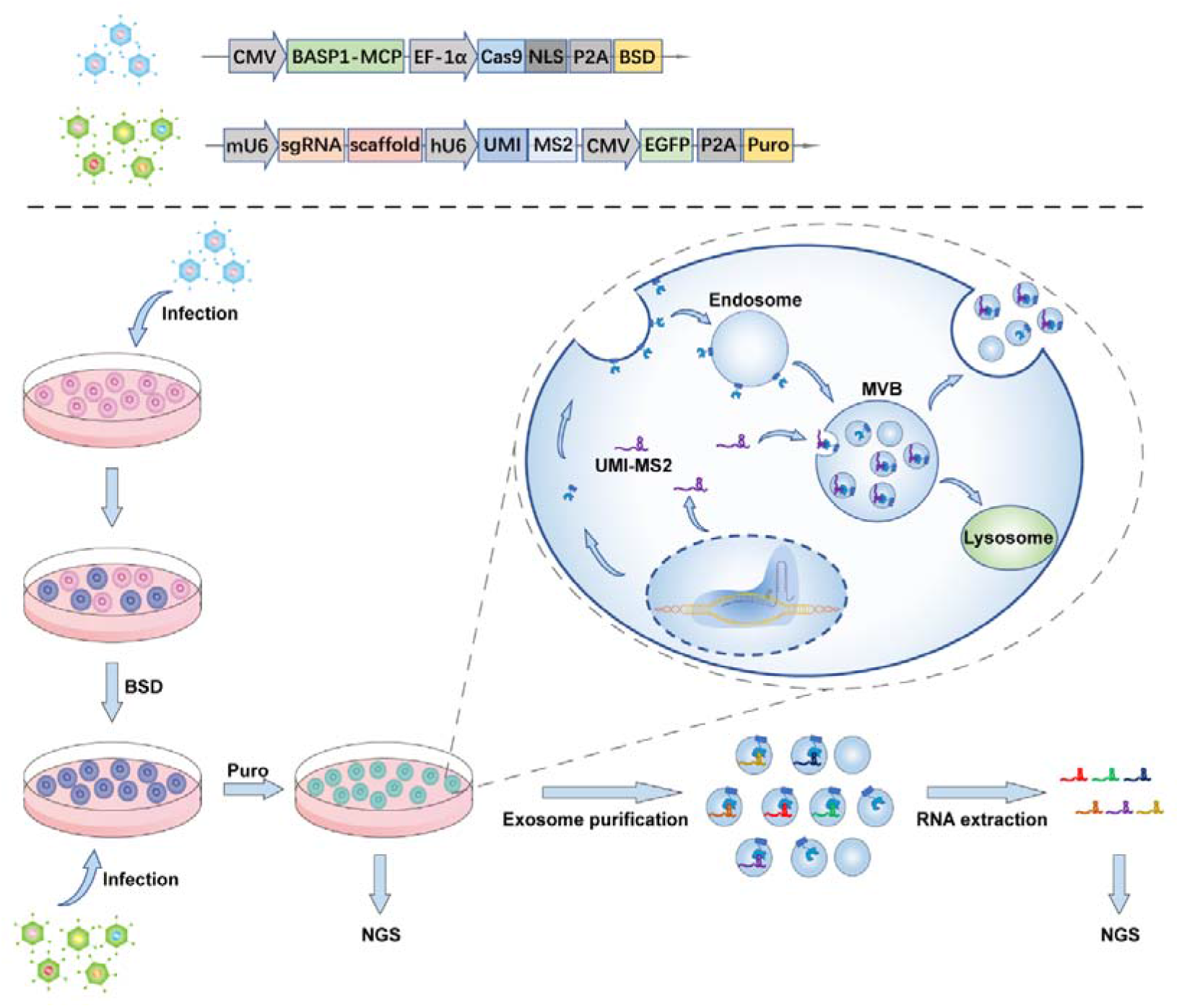
Working principle of the screening system (upper part of the picture). The 293T cells were successively infected by BASP1-MCP-Cas9 and sgRNA-UMI lentivirus (the expression frames of both viruses are shown in the bottom part of the picture). With the 293T cell being stably infected by BASP1-MCP lentivirus, the cell line was infected by sgRNA-UMI lentivirus again. The interaction of two kinds of lentivirus, on the one hand, makes the reduced expression of target genes. On the other hand, exosomes originating from different cells (with different sgRNAs) can be labeled by their corresponding UMIs.

### Functional verification of the screening system’s components by transient transfection

Before we prepared the two kinds of functional lentivirus described in Figure 1, we transient transfected the transfer plasmid to verify the effectiveness of the individual components in the screening system. As exhibited in Figure 2A, the two kinds of sequences were loaded in the plasmids to transfect the 293T cells. For the sgRNA, we chose two target genes, RAB7A and CD276, while the NC (nontargeting) served as control. Based on that, we set seven different groups among these four plasmids. The single transfection of BASP1-MCP-Cas9 plasmid or sgRNA-UMI plasmid was to demonstrate that either plasmid alone in this system cannot work. The BASP1-MCP-Cas9 plasmid was co-transfected with the sgRNA-UMI plasmid to demonstrate that the target gene was knocked out under Cas9 and sgRNA interactions. Meanwhile, the UMI-MS2, corresponding one-to-one with sgRNA, was brought into the exosomes under MCP-MS2 interaction.

**Figure 2.**
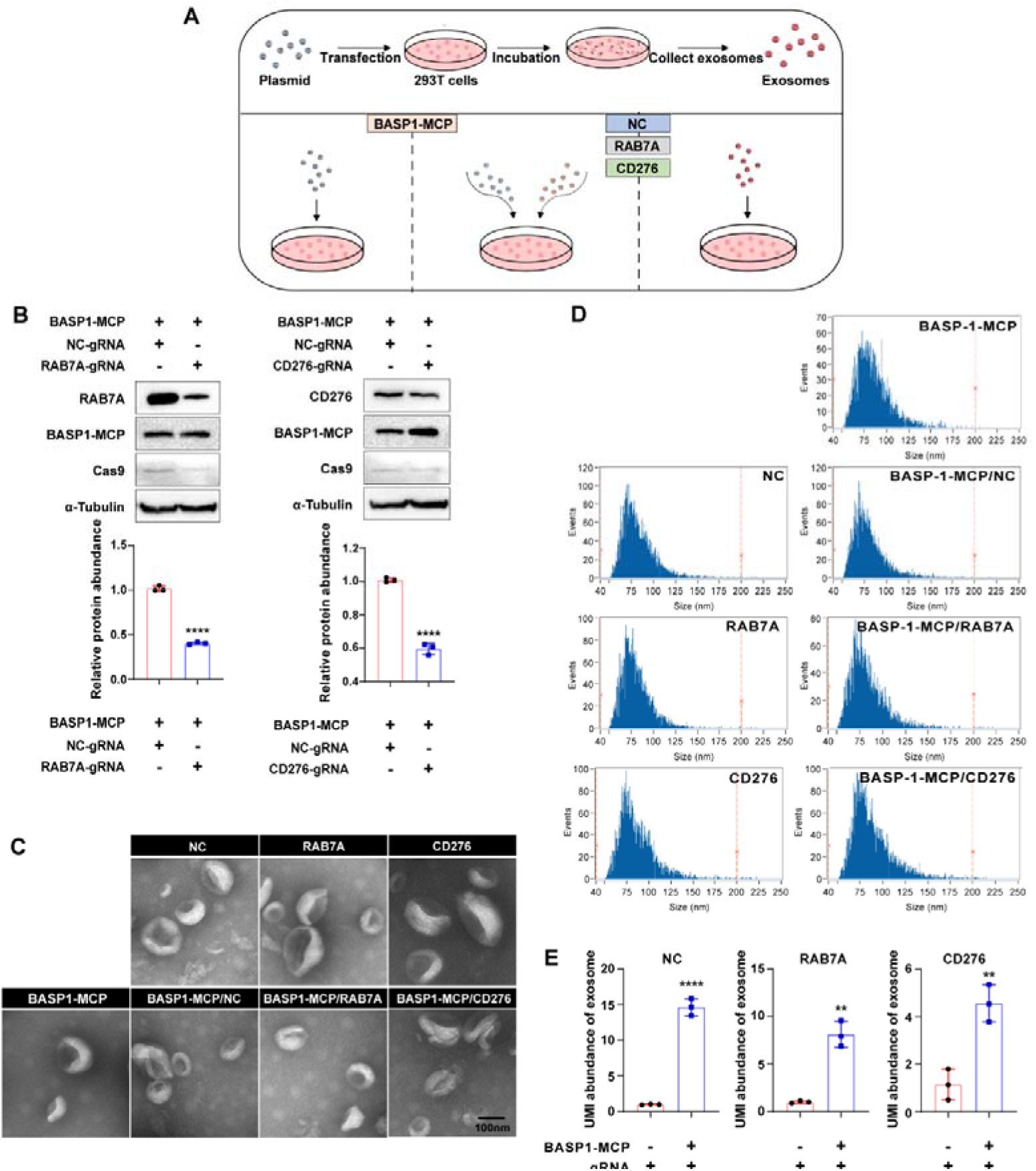
Validation of the screening system by transient transfection. (A) The schematic diagram of the transient transfection system. (B) The expression of RAB7A, CD276, BASP1-MCP, Cas9, and α-tubulin. The relative expression of RAB7A and CD276 were compared to α-tubulin and analyzed by ImageJ. n=3, ****P<0.0001. (C) The morphology of exosomes secreted by 293T cells transfected with different plasmids, which were captured by TEM. Scale bars, 100nm. (D) The diameter of exosomes secreted by 293T cells transfected with different plasmids, which were distributed among 50∼150nm. (E) The content of UMI-MS2 carrying NC, RAB7A, and CD276 sgRNA in the exosomes from the co-transfected group compared to that in the exosomes from 293T cells transfected with sgRNA-UMI plasmid. n=3, **P<0.01, ****P<0.0001.

3×10^6^ cells were collected along with 45ml cell supernatant, respectively. We detected Cas9, BASP1-MCP (flag-tag), RAB7A, and CD276 protein expression levels in each group of cells. Figure 2B (upper) shows that Cas9 and BASP1-MCP proteins were expressed in the cells in all three BASP1-MCP-Cas9/sgRNA-UMI co-transfected groups. Meanwhile, compared with the BASP1-MCP-Cas9/NC group, the RAB7A protein in the BASP1-MCP-Cas9/RAB7A group and the CD276 protein level in the BASP1-MCP-Cas9/CD276 group were significantly lower, respectively (Figure 2B lower). This indicates that Cas9 in this screening system can effectively knockout the gene under its guidance of sgRNA.

To construct the screening system of exosomes-related genes, it is important that the physiological features of exosomes, such as morphology, cannot be affected. Therefore, in order to verify that the components applied in our screening system have no effect on the production of exosomes and their properties, we first examined the morphology and particle size distribution of the exosomes isolated from the supernatant of 293T cells in different groups. From the results, the morphologies of the seven groups of exosomes were basically the same, all showing saucer-like lipid vesicles [17] (Figure 2C). Besides, the diameters of these exosomes were uniformly distributed at about 50-150 nm, which was consistent with the reported results [18] (Figure 2D). Combined with the results of exosomes morphology and particle size distribution, it was shown that these components did not affect the exosomes produced by the cells after transfection, and their physiological characteristics remained almost unchanged. Then we compared the levels of UMI-MS2 in the exosomes produced by each group of cells. The results showed that UMI-MS2 were significantly higher in exosomes from the BASP1-MCP-Cas9/sgRNA-UMI co-transfected group than the sgRNA-UMI single transfected group (Figure 2E). This suggests that UMI-MS2 can be actively loaded into exosomes with the help of BASP1-MCP in exosomes.

### Functional verification of the screening system’s components by stable transfection

Based on the above validation, we prepared the two kinds of functional lentivirus mentioned in Figure 1, to verify whether all the components worked as expected. Using lentivirus as the delivery vehicle, the experimental setups were almost identical to Figure 2A, except for the mode of infection. 293T cells were first infected by the lentivirus containing the BASP1-MCP-Cas9 component. A BASP1-MCP-Cas9 monoclonal cell line was selected after 10 days of BSD resistance screening to eliminate systematic bias caused by cell heterogeneity (Figure 3A). As shown in Figure 3B, the cell line was re-infected by lentivirus carrying NC-sgRNA, RAB7A-sgRNA, and CD276-sgRNA, respectively. Then, the EGFP expression in the cells in different groups (sgRNA-UMI lentiviral vectors containing EGFP gene) was analyzed. The results showed that the sgRNA-positive groups all emitted green fluorescence, while the flow cytometry analysis reflected a high rate of sgRNA lentivirus infection of 100% (Figure 3B, 3C). These results indicated that we successfully constructed three double-positive (BASP1-MCP-Cas9/sgRNA-UMI) stable cell lines. To verify whether the target genes were knocked out in these three cell lines, we analyzed the protein expression of RAB7A and CD276 in these cells. Among them, the BASP1-MCP-Cas9 cell line was used as a blank control, and the BASP1-MCP-Cas9/NC-sgRNA cell line was used as a negative control. RAB7A and CD276 expression in the BASP1-MCP-Cas9/NC-sgRNA cell line was almost like the blank group. And the expression of RAB7A and CD276 decreased in BASP1-MCP-Cas9/RAB7A-sgRNA-UMI and BASP1-MCP-Cas9/CD276-sgRNA-UMI cell lines, respectively (Figure 3D (upper)). The results of the grayscale analysis plot (Figure 3D, lower) were consistent with the western blotting results. This indicates that the Cas9 protein of the screening system also had functions in the stable transient cell lines.

**Figure 3.**
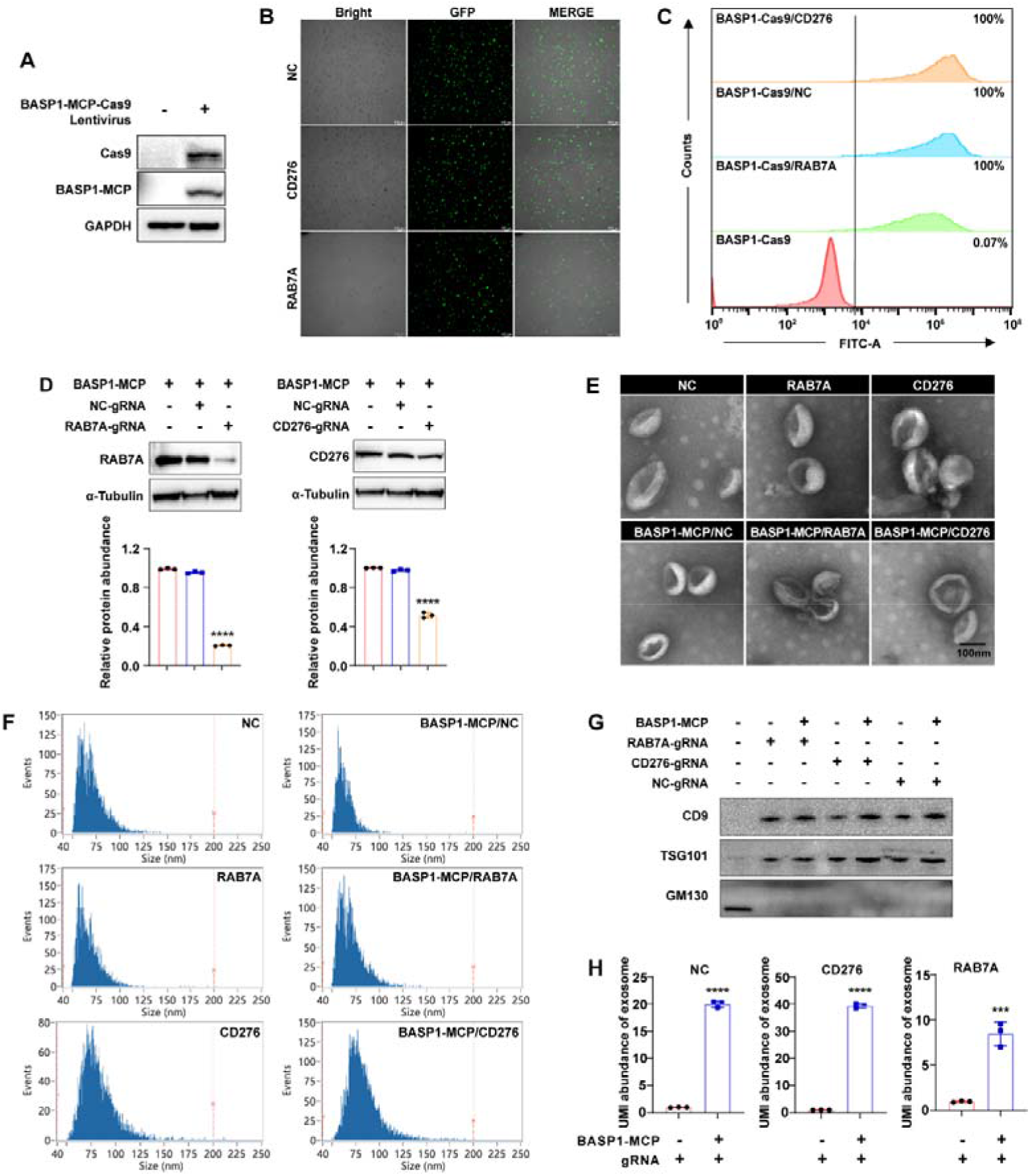
Validation of the screening system by stable transfection. (A) Validation of cas9 and basp1-MCP expression in monoclonal cell lines. (B) The fluorescence intensity of GFP in the 293T cell infected with the BASP1-MCP-Cas9 lentivirus and the lentivirus carrying NC, RAB7A, or CD276 gRNA, which were co-expressed with the UMI-MS2 synergistically. Scale bars, 100nm. (C) FITC positive ratio of cells in different groups was detected by flow analysis. (D) The expression of RAB7A, and CD276 in the 293T cell infected with the BASP1-MCP-Cas9 lentivirus and the lentivirus carrying NC, RAB7A, or CD276 gRNA, which were analyzed through western blotting. The relative expression of RAB7A and CD276 compared to Blank and NC groups, were analyzed by ImageJ. n=3, *P <0.0001. (E) The morphology of exosomes secreted by 293T cells transfected with different plasmids, which were captured by TEM. Scale bars, 100nm. (F) The diameter of exosomes secreted by 293T cells transfected with different plasmids, which were distributed among 50∼150nm. (G) Detection of exosomes surface positive and negative markers. (H) The content of UMI-MS2 carrying NC, RAB7A, and CD276 sgRNA in the exosomes from the re-infected group compared to that in the exosomes from 293T cells only infected with sgRNA-UMI lentivirus. n=3, ***P<0.001,****P<0.0001.

Then, we identified the exosomes isolated from each group of cell culture supernatants in terms of morphology, particle size distribution, and surface markers. All groups of exosomes showed a typical saucer-like structure with no significant differences (Figure 3E). Meanwhile, the particle diameters of exosomes in each group were concentrated in the 50-150 nm range, consistent with the relevant reports (Figure 3F). Exosomes from all groups expressed the exosomes-positive markers, CD9 and TSG101, while there was no expression of the exosomes-negative surface markers GM130 (Figure 3G). Taken together, these exosomes identifications indicate that we successfully isolated the exosomes produced by each group of cells and that there were no significant changes in the exosomes generated by the cells caused by lentiviral infection. In addition, the loading of UMI-MS2 in the BASP1-MCP-Cas9/sgRNA double-positive group was significantly higher than in the sgRNA single-positive group (Figure 3H). The results were consistent with the effects of transient transfection (Figure 2E). This indicates that in the stably transfected cell line, UMI-MS2 can also be actively loaded into exosomes with the help of BASP1-MCP recognizing MS2 in exosomes.

### Screening system validation when each cell was infected with a single sgRNA

Before infecting BASP1-MCP-Cas9 monoclonal cells with the sgRNA-UMI lentiviral library, it is necessary to ensure the incubation time required for desired knockout efficiency. We mimicked the lentiviral library with RAB7A-sgRNA-UMI inserted lentivirus, and infected BASP1-MCP-Cas9 positive monoclonal at low multiplicity of infection (∼0.3 MOI) as described[19, 20]. We selected the sgRNA-transduced cells with puromycin after 72h of incubation. Fluorescent photographs were taken at regular intervals, and the results showed that the proportion of GFP-positive cells increased with the extension of the resistance screening (Figure 4A). We examined the expression of RAB7A in the cells at different time points, and Figure 4B (upper) shows that the expression of RAB7A in the cells decreased gradually with the prolongation of the culture time. The grayscale analysis plot shows the relative expression of RAB7A (Figure 4B, lower).

**Figure 4.**
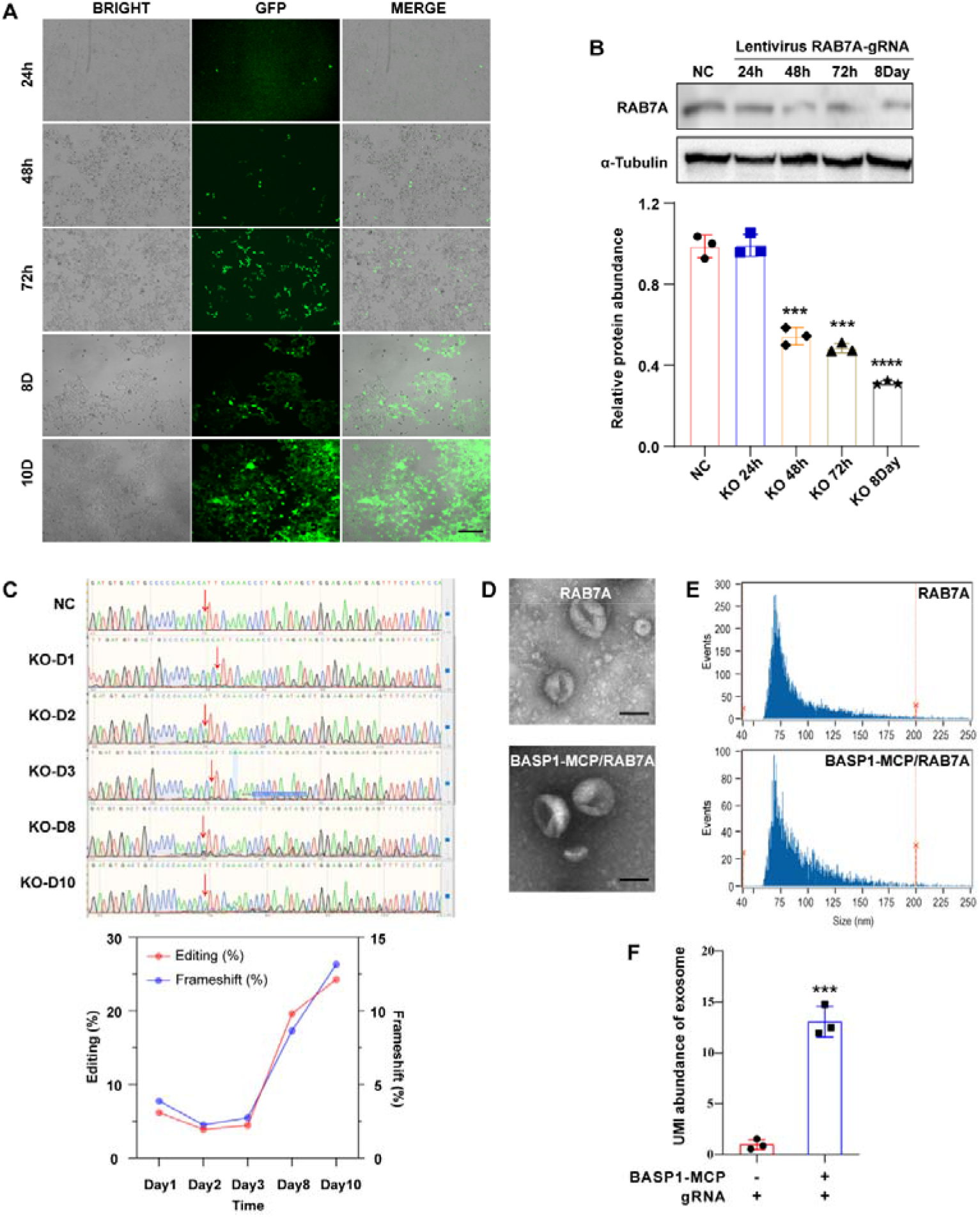
Screening system validation when each cell was infected with a single sgRNA. (A) Expression of EGFP at different time points after each BASP1-MCP-Cas9 cell was infected by a single RAB7A-sgRNA lentivirus. Scale bars, 100nm. (B) RAB7A expression in cells at different times after RAB7-sgRNA lentivirus infection. Their relative expression compared to NC groups, was analyzed by ImageJ. n=3, ***P<0.001 , ****P<0.0001. (C) Upper: Sanger sequencing results of RAB7-sgRNA targeting regions of cells at different times. The DNA sequence of the 293T cell only infected by the RAB7-sgRNA lentivirus served as control. Base mutations occur wherever the red arrows point, except in the control group. Bottom: The proportion of cellular DNA undergoing gene editing at different times compared to control. (D) The morphology of exosomes originating from different groups of cells, which were captured by TEM. Scale bars, 100nm. (E) The diameter of exosomes originating from different groups of cells, which were measured by NanoFCM. (F) The content of UMI-MS2 of exosomes originating from different groups of cells. n=3, ***P<0.001.

The reduction in RAB7A expression was due to the editing of the RAB7A gene by Cas9 protein under the guidance of sgRNA. In order to explore the time point at which gene editing begins, the DNA of the region targeted by RAB7A-sgRNA from different time points after RAB7A-sgRNA lentivirus infection was verified by Sanger sequencing. The DNA derived from 293T cells infected only with RAB7A-sgRNA lentivirus as a control. In terms of results, the RAB7A-sgRNA targeting region was significantly mutated after 24h of lentivirus infection, and the disorder of DNA sequence increased with the extension of the puromycin resistance culturing time (Figure 4C). The above results indicate that the sgRNA is equally capable of guiding Cas9 for gene editing of target genes while each cell is infected by a single lentivirus. Next, we purified and characterized exosomes produced by single sgRNA lentivirus-infected BASP1-MCP-Cas9 monoclonal cells and 293T cells, respectively (Figure 4D and 4E). The morphology of exosomes originating from the two groups of cells was not significantly different, with a particle size distribution between 50-150 nm. Meanwhile, the loading of UMI-MS2 molecules in BASP1-MCP-Cas9/RAB7A-sgRNA double-positive exosomes was significantly higher than in RAB7A-sgRNA single-positive exosomes. This indicates that MCP/MS2 still works appropriately in the screening system while each cell is infected by a single lentivirus (Figure 4F).

### Systematic validation of the screening library by lentiviral libraries

After confirming that these components worked properly, we designed a lentiviral library consisting of 18 sgRNA-UMIs containing 3 NC-sgRNAs based on previous literature (sgRNAs are listed in Table 1) to verify that the CRISPR-UMI-based screening system worked well. First, the appropriate lentivirus quantity was determined by adding different amounts of lentivirus for infection, including 0, 0.3, 0.5, and 1MOI. The FITC-positive rates of the cells were all less than 30%. To make the validation results more reliable, we chose a relatively low MOI (MOI=0.5). Based on that, we performed screening of selected experimental groups using puromycin (Figure 5A). When the percentage of cells positive for FITC exceeded 99%, we re-inoculated and cultured these cells, and the cell culture supernatants were collected after 72h for exosomes isolation and purification. The morphology of exosomes was not affected, which showed a consistent saucer-like vesicle structure (Figure 5B). The diameter distribution also exhibited a similar mode to that detected in the above results (Figure 2C), mainly concentrated between 50-150nm (Figure 5C). Inherently, the proteins of CD81 and TSG101 were highly expressed in the exosomes compared to that in the 293T cells and the expression of GM130 was negative (Figure 5D). And, the HPLC analysis result also showed that the purity of exosomes reached 100% (Figure 5E).

**Table 1.** Genes associated with exosomes biogenesis.

| Gene | Roles | Pathways | Refs |
| --- | --- | --- | --- |
| RALA | + | Syndecan-syntenin-ALIX | [21] |
| RAB7A | - | Syndecan-syntenin-ALIX | [22] |
| VPS4A | / | / | [23] |
| RAB27A | + | SMPD3 pathway | [24] |
| RAB11A | / | / | [25] |
| ALIX | + | Syndecan-syntenin-ALIX | [23] |
| RAB27B | + | ESCRT independent | [24] |
| SDCBP | + | Syndecan-syntenin-ALIX | [23] |
| TSG101 | + | ESCRT independent | [26] |
| Hrs | + | ESCRT independent | [26] |
| RAB35 | + | ESCRT independent | [27] |
| Vps25 | + | / | [28] |
| ATG5 | + | / | [29] |
| SMPD3 | + | SMPD3 pathway | [30] |
| STAM1 | + | ESCRT independent | [26] |

**Figure 5.**
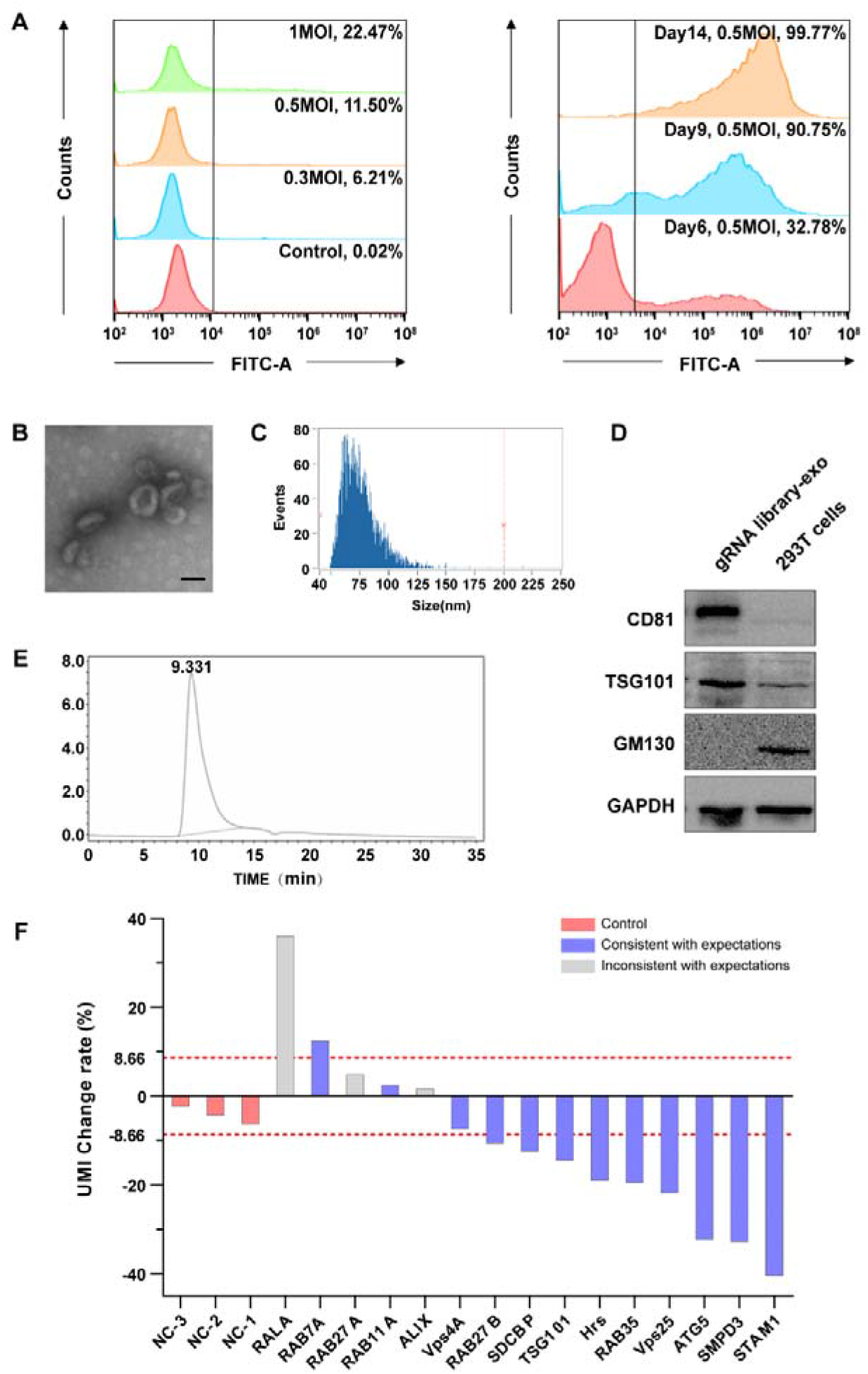
Systematic verification of the screening system by the lentiviral library. (A) The flow cytometry analysis of lentivirus infection rates at different MOI (Left). As the puromycin screen progressed, the percentage of FITC-positive cells increased (Right). (B) The morphology of exosomes collected from infected cells was captured by TEM. Scale bars, 100nm. (C) The diameter of exosomes collected from infected cells was analyzed by NanoFCM. (D) The expression of CD81, TSG101, and GM130 in exosomes was collected from infected cells, which were compared to that in the 293T cells. The expression of GAPDH served as control. (E) The purity analysis of exosomes collected from infected cells, which were close to 100%. (F) The change rate of UMI-MS2 in infected cells ranged between -40% and 40%. Purple columns indicate that the trend is consistent with expectations, and gray indicates that the trend is inconsistent with expectations.

Further, as the results presented, the UMI-MS2 change rates were distributed between -40% and 40% (Figure 5F). Combined with the working principle of the system reflected in Figure 1 and the validation of the efficiency of the screening system, the results showed 11 UMIs with a rate of change greater than the threshold (2 times the absolute mean of the rate of change for NC1, NC2, and NC3). Ten of the 11 UMIs were in line with the expected trend of change and one (RALA-UMI) was the opposite of the expected trend of change. Of the remaining 4 UMIs that did not exceed the threshold, 2 UMIs corresponded to genes for which previous reports indicated that their knockout did not affect exosomes production. It suggests that there are other pathways for exosomes biogenesis that are currently unknown to us. This data stated that the CRISPR-UMI-based screening system was successfully onstructed, and 80% (12/15) of the genes in this lentiviral library had effect on exosomes biogenesis consistent with previous reports.

## Discussion

In this study, we developed a screening system that enables both negative and positive selection screening genes related to exosomes biogenesis including production and secretion. We validated the effectiveness of the system for screening a small library consisting of 15 reported exosomes biogenesis-related and 3 negative controls. 80% of the genes relative to exosomes biogenesis were effectively screened.

As for the enhancement of exosomes production, the existing means are mainly through physical, chemical, and biological methods to stimulate the cells to generate more vesicles. Physical processes usually require special equipment, while chemical methods may cause chemical penetration and residue[31]. All these pose great challenges to the production and purification process of exosomes. Systematically directed screening of genes related to exosomes biogenesis, followed by genetic engineering to endow cells with new vitality and construct cells suitable for industrial production, is an effective way to improve the production of exosomes, which is also in line with the current concept of synthetic biology. This helps the exosomes production process to match the existing GMP production process. Therefore, an effective screening system is necessary. One of the ingenious designs in this CRISPR-UMI-based lentivirus screening system is that it brings UMI-MS2 into exosomes using a fusion protein of an exosomes localization protein and RNA binding protein, which effectively minimizes the background error caused by the low UMI-MS2 loadings of the exosomes naturally[32]. The other ingenious design of this screening system is the precise one-to-one correspondence between sgRNA and UMI-MS2[33], which avoids mismatching of UMI-MS2 and sgRNA in bioinformatics analysis after high-throughput sequencing and consequent misidentification of genes.

This study validated our screening system with a sgRNA-UMI library containing 18 kinds of sgRNAs in a BASP1-MCP-Cas9 positive monoclonal cell line. The results demonstrated the feasibility and efficiency of this screening system, but there is still room for improvement in the accuracy of the screening. This may be due to the limited gene knockdown effects of the individual sgRNAs in this 18 sgRNA-UMIs lentiviral library, resulting in weak changes in exosomes production and secretion, not sufficiently reflecting the role of the gene in exosomes biogenesis. Meanwhile, a genome-wide sgRNA-UMI lentiviral library is being screened; in which each gene has three sgRNAs corresponding to it, ensuring that at least one sgRNA is able to guide Cas9 to efficiently knockout genes. Using a genome-wide library screen may significantly improve the accuracy of this screening system. The results of the genome-wide screen will be presented in the next version of the manuscript.

This system can serve both industrial cell line construction and scientific research. For example, we can mix exosomes containing more than 58,000 kinds of UMI (corresponding to sgRNAs) produced in this system co-incubating them with recipient cells to screen the gene having an impact on the uptake of exosomes. Or exosomes are co-incubated with multiple receptor cells; the uptake preferences of different cells for exosomes can be compared. In addition, when these recipient cells are derived from different tissues or organs, the issue of exosomes targeting can be investigated. Targeted enrichment of exosomes can be achieved by engineering the product cells *in vitro*, which provides the material basis for targeted drug delivery of exosomes.

The screening system we developed is a modular platform that allows the replacement of different scaffold proteins on exosomes (BASP1 used here) fused to MCP, as well as screening in different cell types (HEK293T used here) or replacement of MCP/MS2. Any replacement may have an impact on the screening results. Modifying different modules greatly extends our system and applies it to different studies.

To our best knowledge, no genome-wide screening of genes affecting exosomes production and secretion has been performed in existing studies. Developing our screening system will be essential to systematically understand the mechanisms of exosomes production and secretion. It will also provide data to support future studies and greatly facilitate the progress of exosomes research.

## Conclusions

In this study, a novel gene library used for screening genes related to secretion and production was constructed. In this system, the exosomes can be labeled with a unique barcode (UMI) corresponding to the specific gene when the CRISPR/Cas9 system knocks it out. Both plasmid transient transfection and lentiviral stable transfection results validate the effectiveness of the components of this screening system. On this basis, a lentiviral library containing 18 kinds of sgRNAs was used to validate the feasibility and effectiveness of this screening system. The advantage of this system is that it improves the accuracy of gene identification and avoids the misidentification of genes occurring in the bioinformatics analysis. Additionally, after delivery of the exosomes with different UMI to certain types of cells, analysis of the increase in UMI abundance in cells can help reveal the mechanism involved in exosomes uptake and provide some new ideas for targeted therapy *in vivo*. In conclusion, this screening system can powerfully improve the efficiency and precision of gene screening, which can benefit the research related to exosomes.

## Methods

### Plasmid

For the genome-wide CRISPR/Cas9 screen, the gRNA lentiviral plasmid and library and CRISPR/Cas9 expressing plasmid were obtained from Genewiz, Inc. (Suzhou, China). The BASP1-MCP-Cas9-BSD plasmid was modified by inserting a BASP1-MCP fusion protein on a Cas9 expression plasmid (done by Vector Builder) and performed lentiviral packaging by Vector Builder (Guangzhou, China). The mU6-sgRNA-hU6-UMI-MS2 plasmid was modified by inserting a hU6-UMI-MS2 sequence on a lentiviral gRNA plasmid. RAB7A, CD276, and NC sgRNA plasmids were constructed and performed lentiviral packaging by Genewiz, Inc. (Suzhou, China).

### Lentivirus Infection

The HEK293T cells were infected with 10 multiplicities of infection (MOI) of BASP1-MCP-Cas9-BSD lentivirus. Stable expression of BASP1-MCP and Cas9 293T cells were selected with 10ug/ml Blasticidin S (BSD) 72h post lentivector transduction. BSD continuously screened the 293T cells until all the control 293T cells died. Inoculation was performed using a limiting dilution method, monoclonal screening was performed in 96-well plates, and monoclonal cells were verified using Western blotting. BASP1-MCP-Cas9-BSD monoclonal cells were infected with RAB7A, CD276, and NC sgRNA lentiviral particles at 10 MOI, respectively. Cells stably expressing RAB7A, CD276, and NC sgRNA were screened using 1ug/ml Puromycin (Puro). The sgRNA plasmid contains EGFP protein, so flow cytometry was used to detect the FITC-positive rate of cells during cell passage until the FITC-positive rate was greater than 99%.

### Flow cytometry

RAB7A, CD276, NC-sgRNA stably transfected cells and BASP1-MCP-Cas9-BSD monoclonal cells were digested with 0.25% trypsin, resuspended in PBS, and 1×10^6^ cells were taken for flow cytometric detection. Data were collected on an LSR II flow cytometer (BD Biosciences) and analyzed with the CytExpert software (Beckman).

### Isolation and Purification of Exosomes

The supernatant without cell debris was ultrafiltrated at 300KD membrane, and the precipitate was collected by ultracentrifugation at 100000g for 2h and resuspended in PBS. Then the suspension was filtered by a 30-80nm pore-size molecular sieve, and then the sample was concentrated by ultracentrifugation at 100000g for 2h.

### Exosomes RNA extraction and RT-qPCR assay of UMI

The total RNA of exosomes was extracted using Qiagen miRNeasy Serum/Plasma Kit (Qiagen, China) and was used as a template to determine the expression of UMI with the indicated primer. The following primer sequences were used: UMI, forward 5’-ACACTCTTTCCCTACACGACGCT-3’, and reverse 5’-GACTGGAGTTCAGACGTGTGCT-3’, using the 2× SYBR Green Mix (AG, China) in qPCR. The cycling parameters were as follows: a PCR reaction was carried out on 50 ng of cDNA samples using 0.2μmol/L of each primer, and 10μL 2×SYBR Green Mix. The following conditions were used: 95 °C for 30 s, 40 cycles at 95 °C for 5 s, and 60 °C for 30 s in a Thermo Fisher QuantStudio 5.

### Characterization of the exosomes morphology

The morphology of exosomes was characterized by transmission electron microscopy (TEM). First, the exosomes (100μg/mL) were fixed by mixing with an equal volume of 4% (w/v) paraformaldehyde at room temperature for 15 minutes. Next, the sample was loaded on a formvar-carbon-coated TEM grid and kept at room temperature for 3 minutes. Then, the grid was negative staining with 10μL of uranyl oxalate solution (4% uranyl acetate, 0.0075mol/L oxalic acids, pH=7). Images were collected using a Hitachi 7800 transmission electron microscope operated at 120 KV and 50,000 times magnification.

### Particle size distribution of exosomes

The absolute number and size distributions of exosomes were determined by the NanoFCM instrument (NanoFCM Inc., Xiamen, China)

### Statistical analysis

Statistical analysis was performed with GraphPad Prism 9.3 software. Data are presented as mean ± SD of triplicates unless otherwise indicated. Statistical analysis was performed using a two-tailed Student’s t-test. p < 0.05 was considered statistically significant.

## Notes

### Competing Interest Statement

The authors have declared no competing interest.

## References

1. Pan Bt Fau -Johnstone, R.M. and R.M. Johnstone, Fate of the transferrin receptor during maturation of sheep reticulocytes in vitro: selective externalization of the receptor. (0092-8674 (Print)).

2. Zhang, L. and D. Yu, Exosomes in cancer development, metastasis, and immunity. Biochim Biophys Acta Rev Cancer, 2019. 1871(2): p. 455–468.

3. Jeppesen, D.K., et al., Reassessment of Exosome Composition. Cell, 2019. 177(2): p. 428–445 e18.

4. Thakur, A., et al., The mini player with diverse functions: extracellular vesicles in cell biology, disease, and therapeutics. Protein Cell, 2022. 13(9): p. 631–654.

5. Segura, E., et al., ICAM-1 on exosomes from mature dendritic cells is critical for efficient naive T-cell priming. Blood, 2005. 106(1): p. 216–23.

6. Tian, Y., et al., Reduction of choroidal neovascularization via cleavable VEGF antibodies conjugated to exosomes derived from regulatory T cells. Nat Biomed Eng, 2021. 5(9): p. 968–982.

7. Lin, L.Y., et al., Tumor-originated exosomal lncUEGC1 as a circulating biomarker for early-stage gastric cancer. Mol Cancer, 2018. 17(1): p. 84.

8. Castellanos-Rizaldos, E., et al., Exosome-Based Detection of EGFR T790M in Plasma from Non-Small Cell Lung Cancer Patients. Clin Cancer Res, 2018. 24(12): p. 2944–2950.

9. Shi, J., et al., Role of Exosomes in the Progression, Diagnosis, and Treatment of Gliomas. Med Sci Monit, 2020. 26: p. e924023.

10. Cheng, J., et al., Exosomal noncoding RNAs in Glioma: biological functions and potential clinical applications. Mol Cancer, 2020. 19(1): p. 66.

11. Wang, Z., et al., Exosomes decorated with a recombinant SARS-CoV-2 receptor-binding domain as an inhalable COVID-19 vaccine. Nat Biomed Eng, 2022. 6(7): p. 791–805.

12. Jang, S.C., et al., ExoSTING, an extracellular vesicle loaded with STING agonists, promotes tumor immune surveillance. Commun Biol, 2021. 4(1): p. 497.

13. Kamerkar, S., et al., Exosome-mediated genetic reprogramming of tumor-associated macrophages by exoASO-STAT6 leads to potent monotherapy antitumor activity. Sci Adv, 2022. 8(7): p. eabj7002.

14. Tenchov, R., et al., Exosomes horizontal line Nature’s Lipid Nanoparticles, a Rising Star in Drug Delivery and Diagnostics. ACS Nano, 2022. 16(11): p. 17802–17846.

15. Juan, T. and M. Furthauer, Biogenesis and function of ESCRT-dependent extracellular vesicles. Semin Cell Dev Biol, 2018. 74: p. 66–77.

16. Han, Q.F., et al., Exosome biogenesis: machinery, regulation, and therapeutic implications in cancer. Mol Cancer, 2022. 21(1): p. 207.

17. Wang, Y., et al., The Overexpression of TOB1 Induces Autophagy in Gastric Cancer Cells by Secreting Exosomes. Dis Markers, 2022. 2022: p. 7925097.

18. Arabpour, M., A. Saghazadeh, and N. Rezaei, Anti-inflammatory and M2 macrophage polarization-promoting effect of mesenchymal stem cell-derived exosomes. Int Immunopharmacol, 2021. 97: p. 107823.

19. Shalem, O., et al., Genome-scale CRISPR-Cas9 knockout screening in human cells. Science, 2014. 343(6166): p. 84–87.

20. Chen, S., et al., Genome-wide CRISPR screen in a mouse model of tumor growth and metastasis. Cell, 2015. 160(6): p. 1246–60.

21. Hyenne, V., et al., RAL-1 controls multivesicular body biogenesis and exosome secretion. J Cell Biol, 2015. 211(1): p. 27–37.

22. Fei, X., et al., Neddylation of Coro1a determines the fate of multivesicular bodies and biogenesis of extracellular vesicles. J Extracell Vesicles, 2021. 10(12): p. e12153.

23. Baietti, M.F., et al., Syndecan-syntenin-ALIX regulates the biogenesis of exosomes. Nat Cell Biol, 2012. 14(7): p. 677–85.

24. Ostrowski, M., et al., Rab27a and Rab27b control different steps of the exosome secretion pathway. Nat Cell Biol, 2010. 12(1): p. 19–30; sup pp 1-13.

25. Bai, S., et al., Exocyst controls exosome biogenesis via Rab11a. Mol Ther Nucleic Acids, 2022. 27: p. 535–546.

26. Colombo, M., et al., Analysis of ESCRT functions in exosome biogenesis, composition and secretion highlights the heterogeneity of extracellular vesicles. J Cell Sci, 2013. 126(Pt 24): p. 5553–65.

27. Hsu, C., et al., Regulation of exosome secretion by Rab35 and its GTPase-activating proteins TBC1D10A-C. J Cell Biol, 2010. 189(2): p. 223–32.

28. Ruan, Z., et al., Enterovirus 71 non-structural protein 3A hijacks vacuolar protein sorting 25 to boost exosome biogenesis to facilitate viral replication. Front Microbiol, 2022. 13: p. 1024899.

29. Dias, M.V., et al., PRNP/prion protein regulates the secretion of exosomes modulating CAV1/caveolin-1-suppressed autophagy. Autophagy, 2016. 12(11): p. 2113–2128.

30. Zhang, Z., et al., Micro/nano-textured hierarchical titanium topography promotes exosome biogenesis and secretion to improve osseointegration. J Nanobiotechnology, 2021. 19(1): p. 78.

31. Debbi, L., et al., Boosting extracellular vesicle secretion. Biotechnology Advances, 2022. 59.

32. Dooley, K., et al., A versatile platform for generating engineered extracellular vesicles with defined therapeutic properties. Mol Ther, 2021. 29(5): p. 1729–1743.

33. Michlits, G., et al., CRISPR-UMI: single-cell lineage tracing of pooled CRISPR–Cas9 screens. Nature Methods, 2017. 14(12): p. 1191–1197.

